# Neuropilin-1 Mediates SARS-CoV-2 Infection in Bone Marrow-derived Macrophages

**DOI:** 10.1101/2021.04.14.439793

**Authors:** Junjie Gao, Hong Mei, Jing Sun, Hao Li, Yuege Huang, Yanhong Tang, Linwei Duan, Delin Liu, Qiyang Wang, Youshui Gao, Ke Song, Jincun Zhao, Changqing Zhang, Jia Liu

## Abstract

SARS-CoV-2 infection in human can cause medical complications across various tissues and organs. Despite of the advances to understanding the pathogenesis of SARS-CoV-2, its tissue tropism and interactions with host cells have not been fully understood. Existing clinical data have suggested possible SARS-CoV-2 infection in human skeleton system. In the present study, we found that authentic SARS-CoV-2 could efficiently infect human and mouse bone marrow-derived macrophages (BMMs) and alter the expression of macrophage chemotaxis and osteoclast-related genes. Importantly, in a mouse SARS-CoV-2 infection model that was enabled by the intranasal adenoviral (AdV) delivery of human angiotensin converting enzyme 2 (hACE2), SARS-CoV-2 was found to be present in femoral BMMs as determined by *in situ* immunofluorescence analysis. Using single-cell RNA sequencing (scRNA-Seq), we characterized SARS-CoV-2 infection in BMMs. Importantly, SARS-CoV-2 entry on BMMs appeared to be dependent on the expression of neuropilin-1 (NRP1) rather than the widely recognized receptor ACE2. It was also noted that unlike brain macrophages which displayed aging-dependent NRP1 expression, BMMs from neonatal and aged mice had constant NRP1 expression, making BMMs constantly vulnerable target cells for SARS-CoV-2. Furthermore, it was found that the abolished SARS-CoV-2 entry in BMM-derived osteoclasts was associated with the loss of NRP1 expression during BMM-to-osteoclast differentiation. Collectively, our study has suggested that NRP1 can mediate SARS-CoV-2 infection in BMMs, which precautions the potential impact of SARS-CoV-2 infection on human skeleton system.

## Introduction

As of March 27, 2021, SARS-CoV-2 outbreak has caused more than 100 million infections with 2.7 million deaths^1^. Coronavirus Disease 2019 (COVID-19) patients may develop various clinical manifestations, including severe acute pulmonary disease^2–5^, hepatic dysfunction^3,5^, kidney injury^5^, heart damage^3,5^, gastrointestinal^6^, pancreatic symptoms^7^ and olfactory dysfunction^8^. However, due to the lagged, yet possibly long-lasting effects^9^, the impact of COVID-19 on the skeleton system have not been well understood. Bone is the major reservoir for body calcium and phosphorus^10^. Preliminary clinical data have indicated the occurrence of COVID-19-associated calcium metabolic disorders and osteoporosis^11^. Importantly, severe COVID-19 patients have decreased blood calcium and phosphorus levels, in comparison with moderate COVID-19 patients^12^. These observations suggest that SARS-CoV-2 infection may occur in the skeleton system.

Osteoclasts are one of the major cell types in the bone matrix and can mediate bone resorption. Dysfunction of osteoclasts may result in disturbed bone metabolism including osteoporosis and osteopetrosis^13^, which are characterized with perturbed blood calcium and phosphorus levels^14,15^. We speculate that the observed calcium and phosphorus disorders in COVID-19 patients are attributed to dysregulated osteoclasts. It has been known that osteoclasts originate from the fusion of bone marrow derived macrophage (BMM) precursors in the presence of macrophage colony-stimulating factor (M-CSF) and receptor activator of nuclear factor kappa-B ligand (RANKL)^16^.

Macrophages can sense and respond to the infection of viruses and maintain tissue homeostasis^17^. During the infection of SARS-CoV, dysregulated macrophage response can result in rapid progression of COVID-19^18,19^. In the case of SARS-CoV-2, COVID-19 patient-derived macrophages are found to contain SARS-CoV-2 nucleoproteins^20^. Intriguingly, while macrophages from lymph node subcapsular and splenic marginal zone in COVID-19 patients express SARS-CoV-2 entry receptor ACE2^21^, most tissue resident macrophages from human have little expression of ACE2^20^. Particularly, ACE2 has very low expression in bone marrow cells^22^. These studies have revealed largely undefined interaction networks between SARS-CoV-2 and macrophages.

In this study, we investigated the interactions between SARS-CoV-2 and BMMs *in vitro* and *in vivo* and found that SARS-CoV-2 could efficiently infect BMMs *via* an NRP1-dependent manner. We also examined the altered NRP1 expression profiles during aging and BMM-to-osteoclast differentiation and their correlation with SARS-CoV-2 infection.

## Results

### Authentic SARS-CoV-2 infects bone marrow-derived macrophages (BMMs)

To determine the infectivity of authentic SARS-CoV-2 in BMMs, we analyzed the expression of nucleocapsid protein in control and infected BMMs. Confocal imaging validated the infection of authentic SARS-CoV-2, as shown by the expression of nucleocapsid protein in both human BMMs (hBMMs) (Fig. 1a) and mouse BMMs (mBMMs) (Fig. 1b). We then conducted SMART transcriptomic analysis (Fig. 1c) and found that authentic SARS-CoV-2 genes were significantly expressed in infected mBMMs with nucleocapsid gene bearing the highest expression (Fig. 1d).

**Fig. 1.**
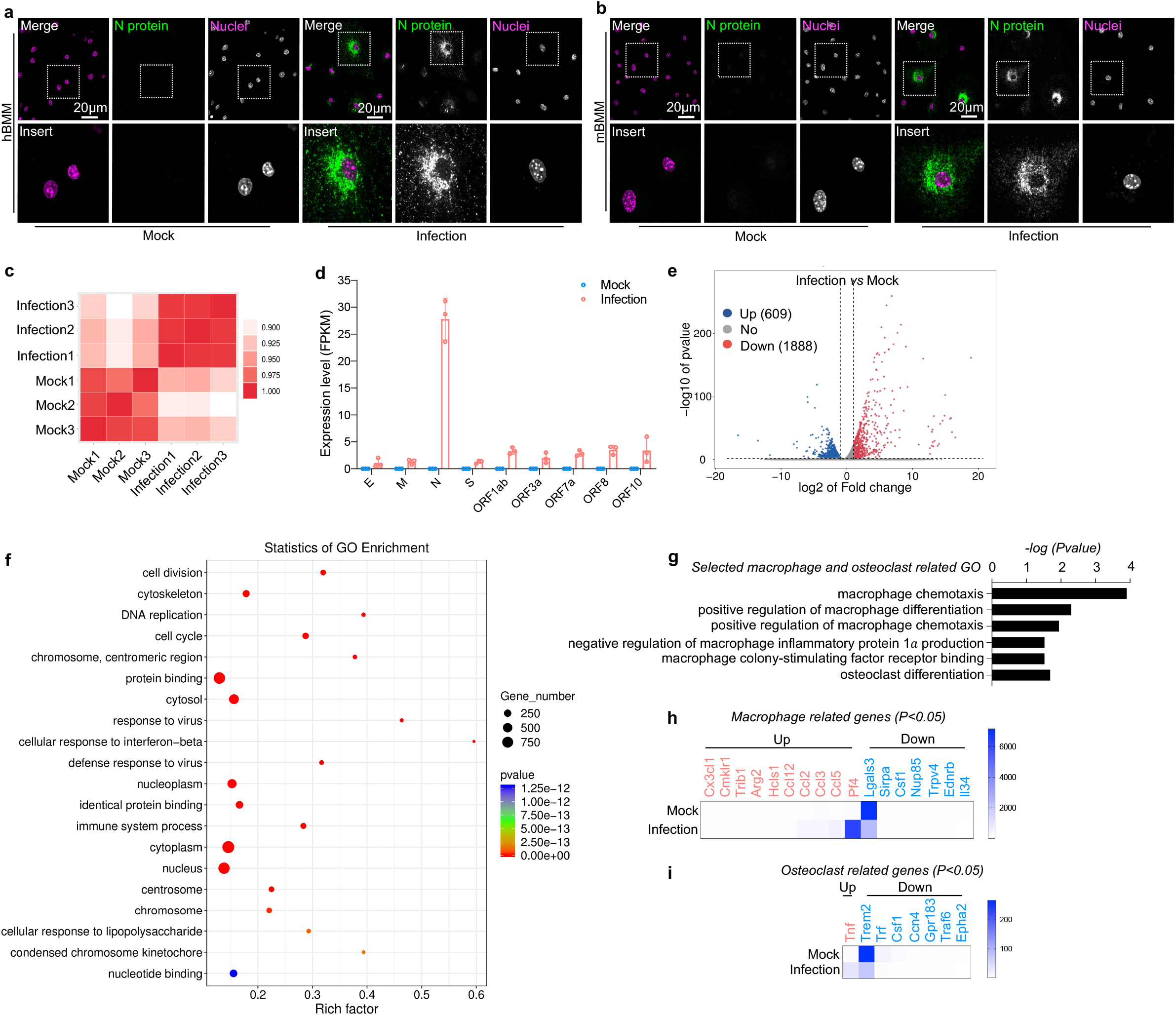
Authentic SARS-CoV-2 infects bone marrow-derived macrophages (BMMs). **a-b**, Confocal images showing the infection of SARS-CoV-2 in human (**a**) and mouse (**b**) BMMs. **c-i**, RNA-Seq analysis of SARS-CoV-2 infection in mouse BMMs. **c**, Heat map showing the correlation analysis of the SMART RNA-Seq results. **d**, Analysis of the expression of viral genes. **e**, Volcano plot showing the profile of altered gene expression. **f**, Top 20 enriched gene ontology (GO) terms. **g**, Significantly enriched GO terms on macrophage and osteoclast-related genes. **h**, Altered macrophage-related genes. **i**, Altered osteoclast-related genes.

We next analyzed the effects of authentic SARS-CoV-2 infection on the gene expression profile of mBMMs. It was found that 609 genes were upregulated and 1888 genes downregulated using a cutoff of 2-fold and a P value of less than 0.05 (Fig. 1e). Gene ontology (GO) enrichment analysis of significantly regulated genes uncovered items of immune response to viral infection (Fig. 1f). Notably, macrophage chemotaxis/differentiation and osteoclast differentiation-related genes were significantly regulated (Fig. 1g), among which Pf4 and Lgals3 had greatest difference in macrophage related genes (Fig. 1h) and Trem2 in osteoclast related gene (Fig. 1i). These results demonstrated that authentic SARS-CoV-2 infection had a global impact on the gene expression profile of macrophages and may perturb the differentiation towards osteoclasts. Nevertheless, we found that authentic SARS-CoV-2 infection in BMMs yielded no detectable productive virus, which is similar to the results of SARS-CoV infection in macrophages^23,24^.

### SARS-CoV-2 infects femoral BMMs in mouse model

As there is currently no established SARS-CoV-2 infection model for the skeleton system, we employed a previously established mouse model for SARS-CoV-2 infection^25^. In this model, adenovirus carrying human ACE2 (hACE2) was intranasally administrated to BALB/c mice at 5 days prior to SARS-CoV-2 infection. Once hACE2 expression in mouse lung was confirmed, SARS-CoV-2 was intranasally administrated and incubated for 2 days (Fig. 2a). Femoral proximal end, midshaft and distal end segments were collected and processed for *in situ* immunofluorescence analysis (Fig. 2a). It was found that SARS-CoV-2 nucleocapsid protein was consistently co-localized with F4/80, the major marker of macrophages^26^, across different segments of femurs (Fig. 2b-d). These results provided critical evidence for the *in vivo* infection of authentic SARS-CoV-2 in BMMs.

**Fig. 2.**
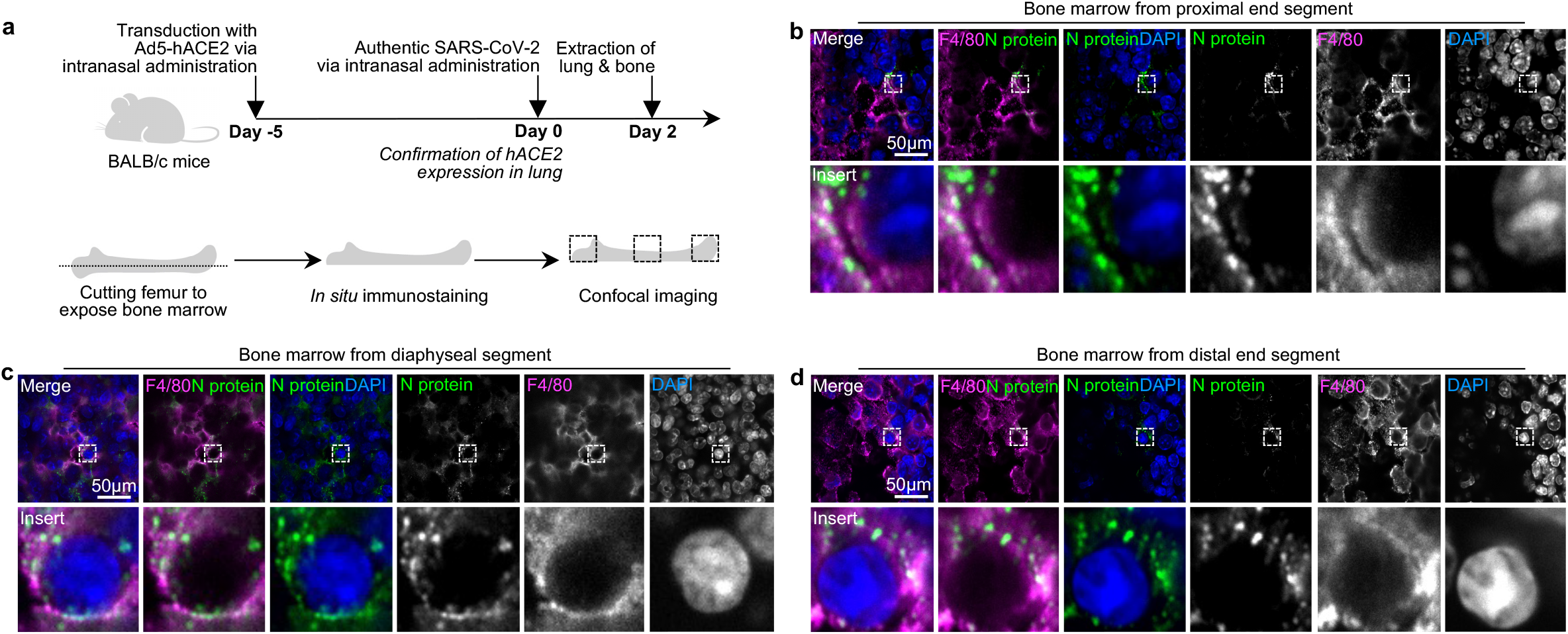
*In situ* immunofluorescence analysis of the *in vivo* infection of authentic SARS-CoV-2 in BMMs of Ad5-hACE2 transduced BALB/c mice. **a**, Schematic illustration of experimental procedures. **b-d**, Confocal images showing the infection of SARS-CoV-2 in BMMs from different femoral segments. SARS-CoV-2 nucleocapsid protein (N protein) and macrophage major marker F4/80 are immunostained.

### SARS-CoV-2 pseudovirus infects BMMs

A key step during SARS-CoV-2 infection in BMMs is virus entry. To dissect essential host factors during this process, we established a lentivirus-based pseudovirus containing SARS-CoV-2 spike protein, where tdTomato was included as the transgene for determination of virus entry (Fig. 3a). It was found that SARS-CoV-2 pseudovirus could efficiently infect *in vitro* cultured hBMMs and mBMMs (Fig. 3b), as determined by the tdTomato expression using RT-qPCR analysis (Fig. 3c-d).

**Fig. 3.**
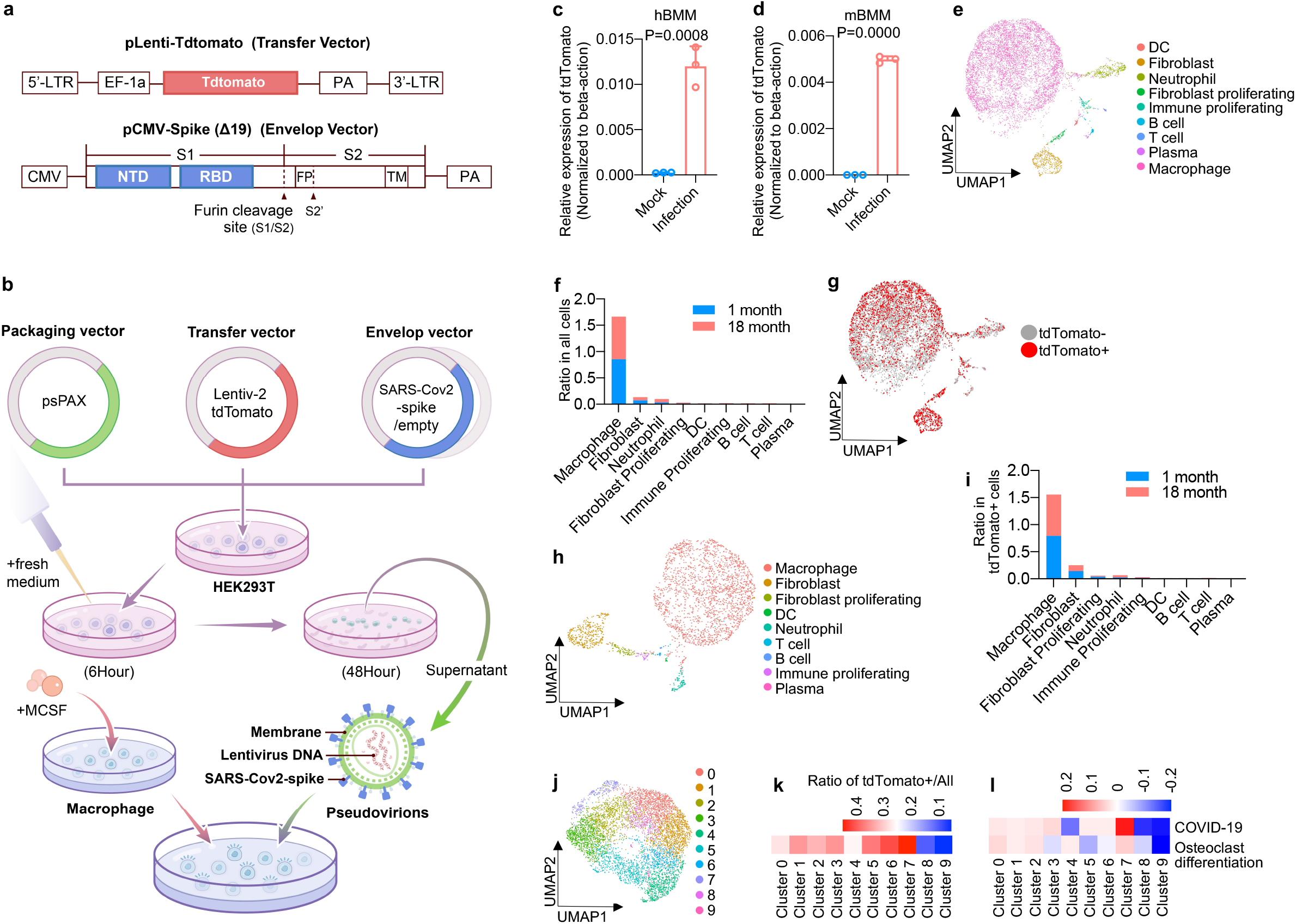
SARS-CoV-2 pseudovirus infects human and mouse BMMs. **a**, Schematic illustration of the design of SARS-CoV-2 pseudovirus. NTD, N terminal domain; RBD, receptor binding domain; PA, poly (A); LTR, long terminal repeats; CMV, cytomegalovirus promoter; EF-1a, elongation factor-1a promoter. **b**, Flowchart showing the procedures of SARS-CoV-2 pseudovirus infection in BMMs. **c-d**, RT-qPCR quantification of SARS-CoV-2 pseudovirus infection in cultured human (**c**) and mouse (**d**) BMMs. **e-l**, Single-cell transcriptome analysis of SARS-CoV-2 pseudovirus infection in cultured BMMs that are derived from 1-month and 18-month mice. **e**, Overview of cell clusters in integrated cell population. **f**, Analysis of cell type ratio, showing that macrophages are the predominant population. **g**, Analysis of SARS-CoV-2 pseudovirus-infected and -uninfected cells, determined by tdTomato transgene expression. **h-i**, Re-cluster of SARS-CoV-2 pseudovirus-infected cells (**h**) and cell number quantitation (**i**), showing that macrophages are the predominant population in SARS-CoV-2 pseudovirus-infected cells. **j-l**, Re-cluster of macrophages. **j**, Cell clusters and analysis of SARS-CoV-2 pseudovirus-infected and -uninfected cells, determined by tdTomato transgene expression. **k**, Cluster 7 shows the highest efficiency of SARS-CoV-2 pseudovirus infection. **l**, Cluster 7 exhibits highest correlation with COVID-19 and osteoclast differentiation.

Next, we collected pseudovirus-infected mBMM that were isolated from the bone marrow of 1− and 18-month mice for single-cell RNA sequencing (scRNA-Seq) analysis. We classified cell groups according to the expression marker genes, and found that macrophages were the major infected cell type (Fig. 3e-f, Extended Data Fig. 1a-c). By re-clustering tdTomato-positive cells, we found that macrophages, neutrophils, proliferating cells, fibroblasts, monocytes, DCs, T cells and B cells were all permissive to the pseudovirus (Fig. 3g-i, Extended Data Fig. 1d-f). Furthermore, we found that pseudovirus-infected macrophages could be grouped into 10 clusters (Fig. 3j, Extended Data Fig. 1g-i) and that cluster 7 had the highest infection rate (Fig. 3k). In consistency with the high infection rate, cluster 7 was found to be highly correlated with COVID-19 and osteoclast differentiation (Fig. 3l, Extended Data Fig. 1j). These results uncovered a possible link between SARS-CoV-2 infection in BMMs and COVID-19 or osteoclast differentiation.

### NRP1 mediates SARS-CoV-2 pseudovirus infection in mBMMs

Although ACE2 is a widely recognized entry receptor for SARS-CoV-2, we found in this study that ACE2 had little expression in *in vivo* isolated mBMMs (Fig. 4a-b, Extended Data Fig. 2a-c) or in *in vitro* cultured hBMMs or mBMMs (Fig. 4d-e). These results indicated that there might exist other proteins facilitating SARS-CoV-2 entry in BMMs. Intriguingly, we found that a previously reported SARS-CoV-2 receptor NRP1 was highly expressed in both *in vitro* cultured BMMs and *in vivo* isolated BMMs (Fig. 4d-e). Sequence alignment and structural analysis showed that human (PDB ID: 7JJC) and mouse NRP1 (PDB ID: 4GZ9) shared remarkable similarity, supporting the consistent results of SARS-CoV-2 infection in hBMMs and mBMMs (Fig. 4f-g). Particularly, the CendR motif from the S1 subunit of SARS-CoV-2 spike protein^27,28^ could be well docked into the b1 domain^27^ of the superimposed NRP1 structures (Fig. 4f).

**Fig. 4.**
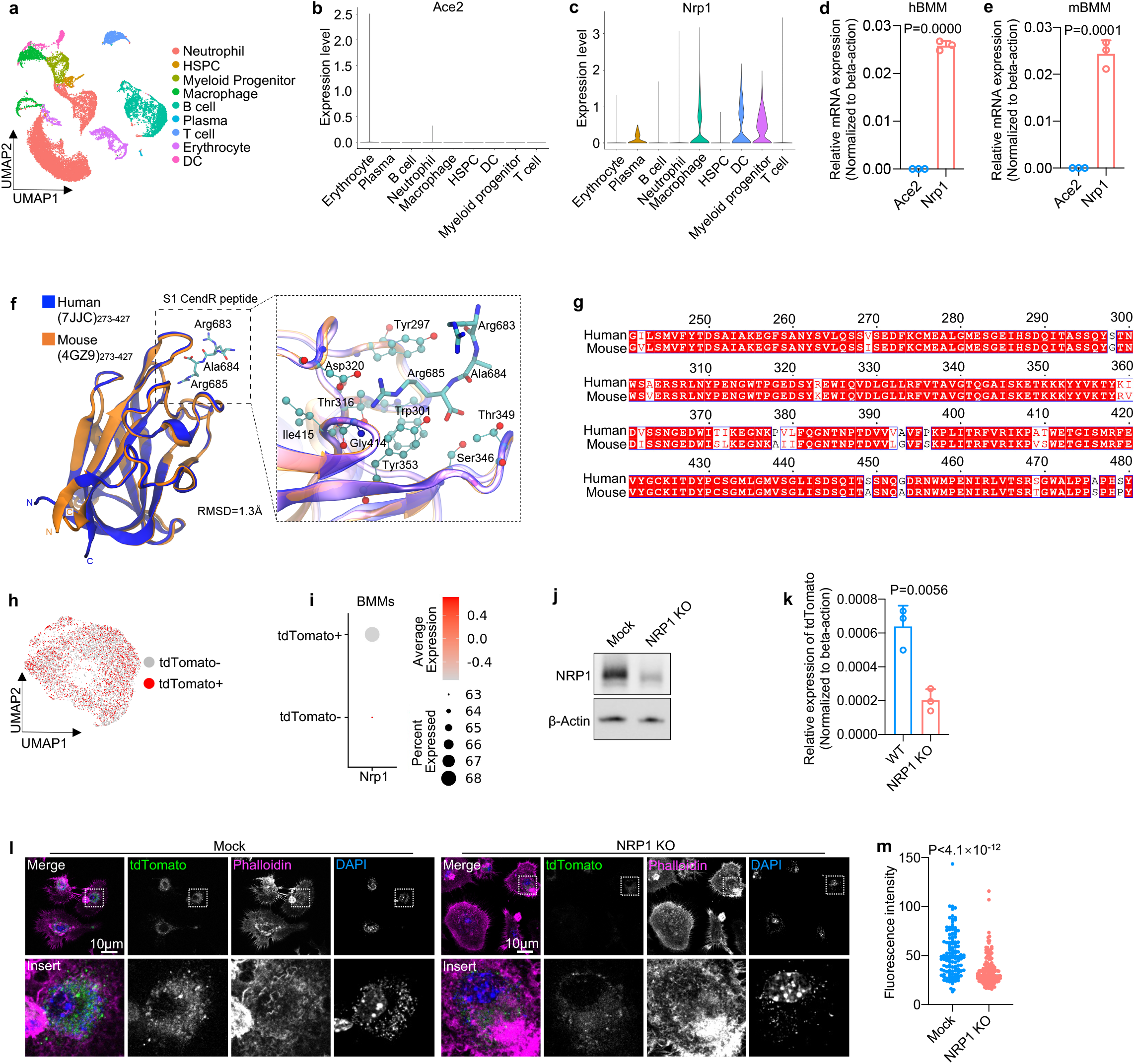
NRP1 facilitates SARS-CoV-2 pseudovirus infection in mBMMs. **a-c**, Single-cell transcriptome analysis of BMMs directly isolated from 1−, 6− or 20-month mice. **a**, Overview of cell clusters in the integrated cell population. Violin plot showing the expression of ACE2 (**b**) and NRP1 (**c**) in each cell type. **d-e**, RT-qPCR quantification of ACE2 and NRP1 expression in cultured hBMMs (**d**) and mBMMs (**e**). **f**, Human NRP1b1-S1 CendR peptide complex superposed with mouse NRP1b1-S1 CendR peptide complex (PDB ID:7JJC and 4GZ9). Binding peptide is shown in stick representation. RMSD, root mean square deviation. Enlarged view highlights the binding of S1 CendR peptide. **g**, Amino acid sequence alignment of human and mouse NRP1b1. **h**, UMAP of tdTomato positive and negative mBMMs. **i**, NRP1 expression in tdTomato positive and negative mBMMs. **j**, Western blot confirming NRP1 knockout in cultured mouse BMMs. Mock, non-targeting sgRNA. **k**, SARS-CoV-2 pseudovirus infection in cultured WT and NRP1 KO mBMMs. **l-m**, Confocal images showing SARS-CoV-2 pseudovirus infection in mock and NRP1 KO mBMMs. **l**, Representative images. **m**, Quantification of the infection efficiency. **k** and **m**, The data are represented as mean ± standard deviation. Statistical difference between mock and NRP1 KO cells is determined using two-tailed Student’s *t* test.

Importantly, in pseudovirus-infected mBMMs (Fig. 4h), NRP1 expression was found to be highly correlated with tdTomato expression (Fig. 4i). To further validate the essentiality of NRP1 expression for SARS-CoV-2 pseudovirus infection, we generated NRP1 knockout mBMMs using CRISPR-Cas9 technology (Fig. 4j). RT-qPCR (Fig. 4k) and confocal imaging (Fig. 4l-m) revealed significantly decreased expression of tdTomato in NRP1 knockout mBMMs. Collectively, these results on pseudovirus suggested that NRP1 played an important role in mediating SARS-CoV-2 entry in mBMMs.

### BMMs and brain macrophages have distinct NRP-1 expression profiles

It is known that NRP-1 had important function in central nervous system (CNS)^29^. Recent studies highlighted a possible role of NRP1 in facilitating SARS-CoV-2 spreading from olfactory bulb to brain^30^. In this study we sought to characterize NRP1 expression in both brain macrophages and BMMs. As age-related mortality has been observed in COVID-19 patients^31^, we analyzed macrophages from mice of different ages. It was found that NRP1 expression in brain macrophages was remarkably increased during brain maturation and aging (Fig. 5a-b, Extended Data Fig. 2d-f). By contrast, NRP1 expression in both *in vivo* isolated BMMs and *in vitro* cultured BMMs had minimum difference during aging (Fig. 5c-d). Importantly, the constant NRP1 expression in the BMMs from neonatal and aged mice was consistent with the similar infectivity of SARS-CoV-2 pseudovirus in these two types of BMMs (Fig. 5e). The difference in programmed time courses of NRP1 expression between brain macrophages and BMMs may render BMMs a particularly vulnerable target during the *in vivo* SARS-CoV-2 infection.

**Fig. 5.**
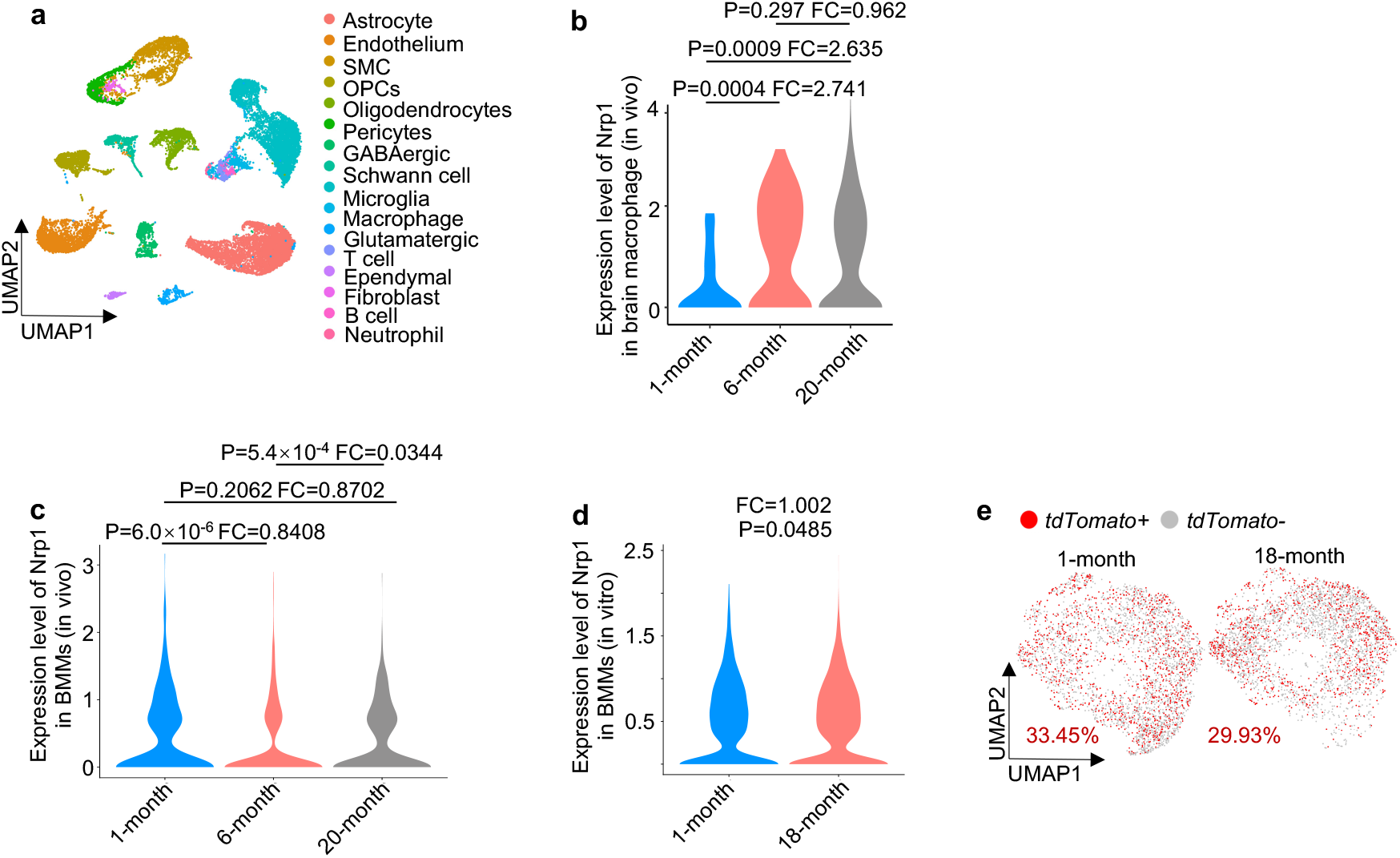
NRP-1 expression profile in BMMs is distinct from that in brain macrophages. **a**, Single-cell transcriptome analysis of brains directly isolated from 1−, 6− or 20-month mice. **b-c**, Violin plot showing the single-cell transcriptome analysis of NRP1 expression in brain (**b**) and bone marrow (**c**) macrophages isolated from 1−, 6− and 20-month mice. Differential gene expression is determined using the function FindMarkers with wilcox rank sum test algorithm under the following criteria: lnFC > 0.25; *P* value < 0.05; min.pct > 0.1. **d**, Violin plot showing the expression of NRP1 in cultured BMMs isolated from 1− or 18-month mice. **e**, Re-clustering of SARS-CoV-2-infected and uninfected cell population in cultured BMMs, determined by tdTomato transgene expression.

### Decreased SARS-CoV-2 pseudovirus infection in mBMM-derived osteoclasts is associated with the loss of NRP1 expression

It has been known that BMMs are one of the main progenitors of osteoclasts^16^. To investigate the infectivity of SARS-CoV-2 on macrophages at different stages during BMM-to-osteoclast differentiation, we infected BMMs with SARS-CoV-2 pseudovirus at the progenitor stage without RANKL stimulation, during differentiation at day 3 after RANKL stimulation, and upon maturation at day 8 after RANKL stimulation, respectively (Fig. 6a and Extended Data Fig. 3). While ACE2 expression was increased by a small degree in mature osteoclasts (Fig. 6b), NRP1 expression was remarkably decreased in differentiating and mature osteoclasts (Fig. 6c-d). Notably, pseudovirus infection appeared to be correlated with the expression of NRP1 during differentiation, where BMMs, the osteoclast progenitors, had higher infection rate than differentiating or mature osteoclasts (Fig. 6e). Additionally, NRP1 expression and the infectivity of pseudovirus had no significant difference between neonatal and aged mouse macrophages (Fig. 6e). Differentiation-related infection was further validated by confocal imaging experiments where higher fluorescence intensity was detected in the perinuclear area of BMMs than in differentiating or mature osteoclasts (Fig. 6f). Interestingly, we found that BMMs expressed Cathepsin B/L but not TMPRSS2 (Extended Data Fig. 4a-c) for the potential priming of SARS-CoV-2^32–34^ and that Cathepsin B/L decreased during aging or BMM-to-osteoclast differentiation (Extended Data Fig. 4d-g). Collectively, these results showed that the reduced infectivity of SARS-CoV-2 during BMM-to-osteoclast differentiation may be a result of decreased NRP1 expression.

**Fig. 6.**
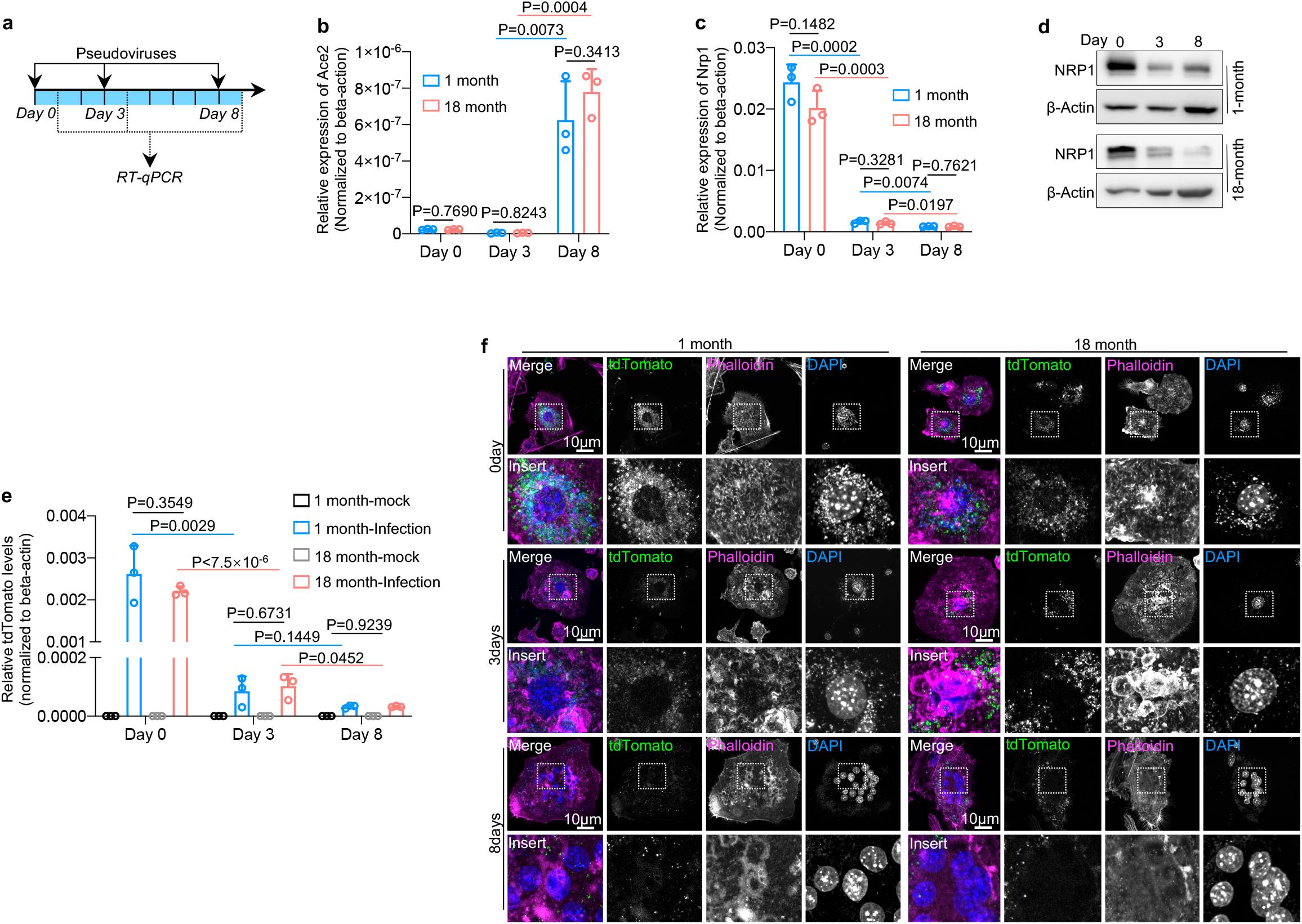
Decreased SARS-CoV-2 infection in mBMM-derived osteoclasts is associated with the loss of NRP1 expression. **a**, Flowchart showing the procedure of BMM-osteoclast differentiation, pseudovirus infection and RT-qPCR quantification. **b-c**, RT-qPCR quantification of ACE2 and NRP1 expression in BMMs (Day 0), deafferenting osteoclasts (Day 3) and mature osteoclasts (Day 8). **d**, Immunoblotting of NRP1 expression in BMMs (Day 0), deafferenting osteoclasts (Day 3) and mature osteoclasts (Day 8). **e**, RT-qPCR quantification of SARS-CoV-2 pseudovirus infection in BMMs (Day 0), differentiating osteoclasts (Day 3) and mature osteoclasts (Day 8), determined by tdTomato transgene expression. **f**, Confocal imaging showing SARS-CoV-2 pseudovirus infection in BMMs (Day 0), deafferenting osteoclasts (Day 3) and mature osteoclasts (Day 8). **b-c** and **e**, The data are shown as mean ± s.d. Statistical difference is determined using two-tailed Student’s *t* test.

## Discussion

In this study, we characterized SARS-CoV-2 infection in BMMs and demonstrated that NRP1 played a critical role during the infection. This work would be the first step toward understanding the causal link between SARS-CoV-2 infection and bone metabolism (Fig. 7). A major challenge to examine the effects of SARS-CoV-2 infection on the skeleton system lies in that there is often lag time between the occurrence of viral infection and detectable disorders of bone metabolism such as osteoporosis or osteopetrosis, though the impact can be long-lasting^9^. Importantly, SARS-CoV-2-associated disorders of bone metabolism is supported by the clinical observation that COVID-19 patients are characterized with disorders of blood calcium and phosphorus^12^.

**Fig. 7.** Proposed model for the effects of NRP-1-mediated SARS-CoV-2 infection on the skeleton system.

Investigation of SARS-CoV-2 infection in the skeleton system is impeded by the lack of an appropriate animal model. Existing SARS-CoV-2 animal models typically rely on the overexpression of hACE2^25,35^. ACE2 is deemed as the entry receptor for SARS-CoV-2. The spike protein of SARS-CoV-2 has a polybasic RRAR (Arg-Arg-Ala-Arg) motif at the boundary of S1 and S2 subunits. During the maturation process, S1 and S2 subunits are cleaved into two polypeptide chains by furin^36^. S1 subunit binds to the RBD domain of ACE2^37^ and S2 mediates membrane fusion following the cleavage by TMPRSS2^33^. In the present study, we employed an SARS-CoV-2 infection model that was enabled by the overexpression of hACE2 in mouse lungs. Thus, the observed SARS-CoV-2 infection in femoral BMMs might be the results of either primary infection originating from injected SARS-CoV-2 or secondary infection originating from replicated viruses that was released from lungs. Nevertheless, establishment of an appropriate animal model can be a critical step toward understanding SARS-CoV-2 infection in the skeleton system.

In recent studies, NRP1 has been proposed as an entry receptor for SARS-CoV-2 in certain cell hosts^27,30^. NRP1 is an important molecule for regulating the maturation of CNS^29^ and may facilitate the spreading of SARS-CoV-2 from olfactory bulb to CNS^30^. NRP1 has been reported to have high expression in infected olfactory epithelial cells of COVID-19 patients^27,30^. Our study has further demonstrated that NRP1 rather than ACE2 played a critical role during SARS-CoV-2 infection in BMMs. Unlike human and mouse ACE2^35^, human and mouse NRP1 has minimum structural difference, justifying the use of mouse BMMs to understand SARS-CoV-2 infection in human BMMs. Intriguingly, NRP1expression was also detected in brain macrophages. Unlike BMMs where NRP1 had relatively constant expression, brain macrophages exhibited aging or maturation-related NRP1 expression. These results may be correlated with the clinical observations that SARS-CoV-2 can infect both young and elder people but cause higher mortality in the latter^38^.

In addition, similar to SARS-CoV^39^, SARS-CoV-2 can take advantage of TMPRSS2 and endosomal cysteine proteases cathepsin B and L in host cells to prime S protein^33^. In TMPRSS2 negative cells, cathepsins B/L seem more predominant in regulating SARS-CoV-2 entry^33^. Interestingly, we found in this study that BMMs had little expression of TMPRSS2 but high expression of cathepsins B/L. Cathepsin B/L expression was also found to be associated with NRP1 expression, exhibiting a decrease during BMM-to-osteoclast differentiation. This and previous studies collectively indicated that the priming process of SARS-CoV-2 spike proteins may be also correlated with the utility of entry receptors.

Finally, although the scRNA-Seq results in our study provided preliminary evidence that SARS-CoV-2 infection may affect BMM-to-osteoclast differentiation, additional experiments are needed to characterize the impact of SARS-CoV-2 infection on bone homeostasis^13^. It is possible that SARS-CoV-2 infection can disrupt the differentiation efficiency or the function of differentiated osteoclasts. Further in-depth studies can help elucidate the impact of SARS-CoV-2 infection on the skeleton system, which may in turn provide insights to the treatment of COVID-19 patients particularly from the perspective of delayed or long-term symptoms.

## Methods

### Cell culture

All experiments including human and animal tissues were approved by the ethics committee of the Shanghai Jiao Tong University Affiliated Shanghai Sixth People’s Hospital. BMMs were obtained from human or mice bone marrows in the presence of human or mice M-CSF (30 ng/ml), and osteoclasts were differentiated from BMMs following the stimulation using human or mouse RANKL (100 ng/ml). BMMs were grown in Minimum Essential Media (alpha-MEM, Thermo) supplemented with 10% fetal bovine serum (FBS, Thermo) and 1% penicillin-streptomycin (Thermo) and maintained at 37 °C in a fully humidified incubator containing 5% CO_2._ HEK293T cells were obtained from the Cell Bank of Shanghai Institutes for Biological Science (SIBS) and were validated by VivaCell Biosciences (Shanghai, China), and were grown in Dulbecco’s modified Eagle’s medium (DMEM, Thermo) supplemented with 10% fetal bovine serum (FBS, Thermo) and 1% penicillin-streptomycin (P/S, Thermo) and maintained at 37 °C in a fully humidified incubator containing 5% CO_2._ Vero E6 cells (derived from African Green monkey kidney) were grown in (DMEM, Thermo) supplemented with 10% FBS (Thermo) and 1% P/S (Thermo). All cells were confirmed by PCR to be free of mycoplasma contamination.

### Authentic SARS-CoV-2 infection

The SARS-CoV-2 strain used in this study was isolated from COVID-19 patients in Guangzhou (Accession numbers: MT123290), and passaged on African green monkey kidney-derived Vero E6 cells. Vero E6 were grown in Dulbecco’s modified Eagle’s medium (DMEM, Gibco, Grand Island, NY) supplemented with 10% FBS. BMMs were infected with SARS-CoV-2 at an MOI of 0.1 or 0.5 for 1-3 days. To analyze the kinetics of viral replication, culture supernatants and cells were harvested at the indicated time points and infectious virus was titrated using a focus forming assay (FFA, see below). Collected cells were lysed using Trizol and fixed using 4% paraformaldehyde (PFA).

### Focus forming assay (FFA)

Vero E6 cells were seeded on to 96-well plates one day before infection. Virus culture was diluted in 1:10 dilution and inoculated on to Vero E6 cells at 37 ? for 1 h. The virus-containing medium was removed and then 1.6% carboxymethylcellulose was added. After 24 h after infection, cells were fixed with 4% paraformaldehyde and permeabilized with 0.2% Triton X-100. Cells were then incubated with a rabbit anti-SARS-CoV nucleocapsid protein polyclonal antibody (Cat. No.: 40143-T62, Sino Biological, Inc. Beijing), followed by an HRP-labelled goat anti-rabbit secondary antibody (Cat. No.: 111-035-144, Jackson ImmunoResearch Laboratories, Inc. West Grove, PA). The foci were visualized by TrueBlue™ Peroxidase Substrate (KPL, Gaithersburg, MD), and counted with an ELISPOT reader (Cellular Technology Ltd. Cleveland, OH). Viral titers were calculated as FFU per ml.

### SARS-CoV-2 infection in Balb/c mouse

Ad5-hACE2 transduced Balb/c mouse model were described previously^25^. Briefly, mice were transduced with 2.5×10^8^ FFU of Ad5-ACE2 intranasally. Five days post transduction, mice were infected with SARS-CoV-2 (1×10^5^ PFU) intranasally. At day 2 post infection, lung and bone were harvested, and fixed in formalin. SARS-CoV-2 infection was conducted in the Biosafety Level 3 (BSL3) Laboratories of Guangzhou Customs District Technology Center. All protocols were approved by the Institutional Animal Care and Use Committees of Guangzhou Medical University.

### Production and infection of SARS-CoV-2 pseudovirus

HEK293T cells were transfected with psPAX, plentiv2-Tdtomato and plasmid that carried SARS-Cov-2 S gene or empty vector using Lipofectamine 3000 (Thermo)^34^. Codon-optimized DNA sequences of S genes were listed in Supplementary Table 1. At 48 h after transfection, supernatant containing pseudovirions was harvested by centrifugation at 2,000 rpm for 10 min and concentrated by Optima XPN-100 Ultracentrifuge (Beckman Coulter, California, USA). For pseudovirus infection, BMMs were seeded on to 6− or 24-well plates (2×10^5^ or 4×10^4^ cells per well) for 24 h, and infected with pseudovirus for 24 h in the presence of polybrene (10 μg/ml, Merck, Darmstadt, Germany). BMMs were washed with PBS for three times at 24 h after infection, and then fixed using 4% PFA for fluorescence detection or lysed using Trizol (Thermo) for RNA extraction.

### Immunofluorescence

Cells were cultured in coverslips (ProSciTech), and fixed in 4% PFA for 20 min. After permeabilizing in 0.1% Triton X-100 in PBS for 5 min, cells were incubated with 3% BSA-PBS for 30 min to block nonspecific antibody binding and then incubated with SARS-CoV-2 nucleocapsid protein antibody (Sino Biological), followed by incubating with Alexa Fluor 488 (Thermo). BMMs were labelled by F4/80 antibody (Abcam), followed by incubating with Cy3-labeled Goat Anti-Rat IgG(H+L) (Beyotime). Nucleus was stained with DAPI (Sigma), and actin cytoskeleton with rhodamine phalloidin (Thermo) or Alexa Fluor 647 phalloidin (Thermo) for 45 min. Immunostained cells were mounted by ProLong Diamond antifade medium (Invitrogen). Images were acquired by Zeiss LSM 710 confocal microscope with EC-Plan-Neofluar 63×oil immersion objective, digital images were acquired by ZEISS ZEN microscope software. Fiji (National Institutes of Health) was employed to analyze and assemble images.

### Immunoblotting

Cell lysate was extracted by incubating in lysis buffer (Sigma) with protease inhibitor (Roche) and a phosphatase inhibitor cocktail (Sigma) for 30 min at 4°C, and diluted with 4×SDS sampling buffer and boiled for 5 min. For each sample, proteins were fractionated on SDS-polyacrylamide gel electrophoresis gel and transferred to a nitrocellulose membrane (Millipore). Membranes were incubated with primary antibodies, including actin JLA20 antibody (Developmental Studies Hybridoma Bank), Neuropilin-1 antibody (Novus Biologicals). Proteins were visualized by enhanced chemiluminescence and autoradiography (FujiFilm LAS-3000/4000 Gel Documentation System).

### Real-time quantitative PCR

Total RNA was purified using TRIzol (Thermo), chloroform (Titan), and isopropanol precipitation. RNA was then reverse transcribed into cDNA by PrimeScript RT reagent Kit with gDNA Eraser (Takara Bio Inc.). Gene mRNA levels were determined using SYBR green dye on Applied Biosystems Q6 Real-Time PCR cycler (Thermo) and specific primers (Supplementary Table 2). All SYBR Green primers were validated with dissociation curves. The expression of genes is normalized to β-actin.

### Generation of CRISPR-Cas9 knockout cells

For pLentiCRISPRsgRNA-v2 construction, pLentivCRISPR-v2 was digested and purified with Esp3I (Thermo) and Gel Extraction kit (Omega). Oligonucleotides encoding the 20 bp sgRNA targeting mouse Nrp1 were synthesized, annealed and ligated to Esp3I-treated pLentiCRISPR-v2 vector using T4 ligase (New England BioLabs) at 37 °C for 1 h. Ligation products were transformed into DH5α *E. coli* (Tsingke), and positive colonies were selected via Sanger sequencing. Plasmid DNA was extracted using NucleoBond Xtra Midi EF kit (Macherey-Nagel). Knockout of target proteins was verified by immunoblotting.

### SMART-Seq

Total RNA of BMMs was extracted using TRIzol (Invitrogen), and then quantified (NanoDrop, Thermo). For SMARTer cDNA synthesis, a modified oligo(dT) primer was employed. When SMARTScribe™ Reverse Transcriptase reaches the 5’ end of the mRNA, the enzyme’s terminal transferase activity adds a few additional nucleotides to the 3’ end of the cDNA. Designed SMARTer Oligonucleotide base-pairs with the non-template nucleotide stretch create an extended template to enable SMARTScribe RT to continue replicating to the end of the oligonucleotide. sscDNA was amplified by LD PCR to get enough cDNA. cDNA was fragmented by dsDNA Fragmentase (NEB, M0348S). For library construction, blunt-end DNA fragments were generated using a combination of fill-in reactions and exonuclease activity. Paired-end sequencing was performed the on an Illumina Novaseq™ 6000 (LC Sciences, USA) following the vendor’s recommended protocol.

### Single-cell RNA Sequencing (scRNA-Seq)

scRNA-Seq experiment was performed by NovelBio Bio-Pharm Technology Co.,Ltd. Bone marrow was flushed and kept in MACS Tissue Storage Solution (Miltenyi Biotec). Samples were sieved through 40μm cell strainers, and centrifuged at 300 g for 5 min. Pelleted cells were suspended in red blood cell lysis buffer (Miltenyi Biotec) for lysing red blood cells. *In vitro* cultured BMMs were washed with 0.04% BSA-PBS, trypsinized and re-suspended. Brain tissues were surgically removed and minced into small pieces (approximately 1 mm^3^) on ice and digested by 200 U/ml Papine (Diamond) and Cysteine-HCL (Sigma). Then samples were sieved through 70 μm cell strainers, centrifuged at 300 g for 10 min, and further cleaned for red blood cells using red blood cell lysis buffer (Miltenyi Biotec). Countstar Fluorescence Cell Analyzer was used for single cells viability assessment, and live cells were further enriched by MACS dead cell removal kit (Miltenyi Biotec).

The scRNA-Seq libraries were generated by 10X Genomics Chromium Controller Instrument and Chromium Single Cell 3’V3.1 Reagent Kits (10X Genomics, Pleasanton, CA). Cells were concentrated to 1000 cells/μL and approximately 8,000 cells were loaded into each channel to generate single-cell Gel Bead-In-Emulsions (GEMs), which results in expected mRNA barcoding of 6,000 single-cells for each sample. After the reverse transcription, GEMs were broken and barcoded-cDNA was purified and amplified. The amplified barcoded cDNA was fragmented, A-tailed, ligated with adaptors and amplified by index PCR. Final libraries were quantified using the Qubit High Sensitivity DNA assay (Thermo) and the size distribution of the libraries were determined using a High Sensitivity DNA chip on a Bioanalyzer 2200 (Agilent). All libraries were sequenced by illumina sequencer (Illumina) on a 150 bp paired-end run.

We applied fastp^40^ for filtering the adaptor sequence and removed the low quality reads. Then the feature-barcode matrices were obtained by aligning reads to the mouse genome (GRCm38 Ensemble: version 92) using CellRanger v3.1.0. Cells contained over 200 expressed genes and mitochondria UMI rate below 20% passed the cell quality filtering, and mitochondria genes were removed in the expression table. Seurat package (version: 2.3.4) was used for cell normalization and regression. Pearson correlation analysis (PCA) was constructed based on the scaled data with top 2,000 high variable genes and top 10 principals were used for UMAP construction. The unsupervised cell cluster results were acquired using graph-based cluster method (resolution = 0.8), and the marker genes were calculated by FindAllMarkers function with wilcox rank sum test algorithm under the following criteria:1. lnFC > 0.25; 2. pvalue<0.05; 3. min.pct>0.1.

## Supporting information

Extended Data Fig. 1 Single cell transcriptome analysis of SARS CoV 2 pseudovirus infection in cultured BMMs from 1 and 18 month mice. a

Extended Data Fig. 2 Single cell transcriptome analysis of BMMs and brain directly isolated from 1 --, 6 and 20 month mice.

Extended Data Fig. 3 RT qPCR quantification of altered gene expression during BMM to osteoclast differentiation.

Extended Data Fig 4 Single cell transcriptome analysis of brain or bone marrow cells directly isolated from 1 --, 6 and 20 month mice.

## Statistical analysis

The data were graphed and statistically analysed using GraphPad Prism 8.0. RT-qPCR data are represented as mean ± standard deviation. Statistical difference is determined using two-tailed Student’s t test. For scRNA-Seq data wilcox rank sum test algorithm was employed under the following criteria:1. lnFC > 0.25; 2. p value<0.05; 3. min.pct>0.1. For SMART-Seq, differentially expressed mRNAs and genes were selected with log2 (fold change) >1 or log2 (fold change) <−1 and with statistical significance (p value < 0.05) by R package.

## Data availability

ScRNA-Seq data have been deposited into GEO repository with accession codes GSE169599, GSE169606 and GSE169608. Additional data that support the findings of this study are available from the corresponding author on request. Source data are provided with this paper.

## Author contributions

J.L., C.Q.Z., J.C.Z. and J.J.G. conceived, designed and supervised the study. J.C.Z. supervised the authentic SARS-CoV-2 work and J.S. performed the infection experiments with assistance from Y.H.T and L.W.D. J.J.G., H.M. and H.L. performed confocal imaging, SARS-CoV-2 pseudovirus, cell culture and differentiation experiments. Y.G.H., D.L.L., Q.Y.W., Y.S.G. and K.S. conducted immunoblotting, RT-QPCR. J.J.G. J.L., H.M., C.Q.Z. and J.C.Z. analysed the data and wrote the manuscript.

## Acknowledgements

We thank Dr. Lichun Jiang and Dr. Wei Wang from High-throughput Screening (HTS) Platform at Shanghai Institute for Advanced Immunochemical Studies (SIAIS), ShanghaiTech University for the support of IF experiments and Biomedical Big Data Platform for the analyses of RNA-Seq data. This work is supported by National Natural Science Foundation of China (81820108020 to C.Q.Z.; 82002339 to J.J.G.; 82025001 to J.C.Z.), National Key R&D Program of China International Collaboration Project (Grant No. 2018YFE0200402 to J.L., 2018YFC1106300 to C.Q.Z.), the emergency grants for prevention and control of SARS-CoV-2 of Ministry of Science and Technology of Guangdong province (2020B1111330001 to J.C.Z.), Zhangjiang National Innovation Demonstration Zone (ZJ2020-ZD-004 to J.L.), China Postdoctoral Science Foundation (2017M621551 to H.M.), ShanghaiTech University Startup Fund (2019F0301-000-01 to J.L.). and Shanghai Sixth People’s Hospital Scientific Research Foundation to J.J.G.

## Extended Data Figures

**Extended Data Fig. 1 Single-cell transcriptome analysis of SARS-CoV-2 pseudovirus infection in cultured BMMs from 1− and 18-month mice. a-c**, Overview of whole cell population. **a**, Sample origin. **b**, Number of transcripts in each cell. **c**, Markers of each identified cell type. **d-f**, Re-clustered tdTomato positive cells. **d**, Sample origin. **e**, Number of transcripts in each cell. **f**, Markers of each identified cell type. **g-j**, Re-clustered macrophage population. **g**, Sample origin. **h**, Number of transcripts in each cell. **i**, Markers of each identified cell type. **j**, QuSAGE analysis of clusters 0-9 in macrophages.

**Extended Data Fig. 2 Single-cell transcriptome analysis of BMMs and brain directly isolated from 1−, 6− and 20-month mice. a and d**, Sample origins. **b and e**, Number of transcripts in each cell. **c and f**, Markers for each identified cell type.

**Extended Data Fig. 3 RT-qPCR quantification of altered gene expression during BMM-to-osteoclast differentiation. a-e**, Up-regulated expression of osteoclastogenesis markers during RANKL stimulation (Day 0, 3 and 8).

**Extended Data Fig. 4 Single-cell transcriptome analysis of brain or bone marrow cells directly isolated from 1−, 6− and 20-month mice. a-b**, Violin plot and RT-qPCR results showing the absence of TMPRSS2 expression in mouse BMMs. **c**, Expression of Ctsl and Ctsb in mouse bone marrow cells. **d-e**, Violin plot showing expression of Ctsb (**d**) and Ctsl (**e**) in BMMs. **f-g**, Decreased expression of Ctsb (**f**) and Ctsl (**g**) during BMM-to-osteoclast differentiation.

---

**Supplementary Table 1.**
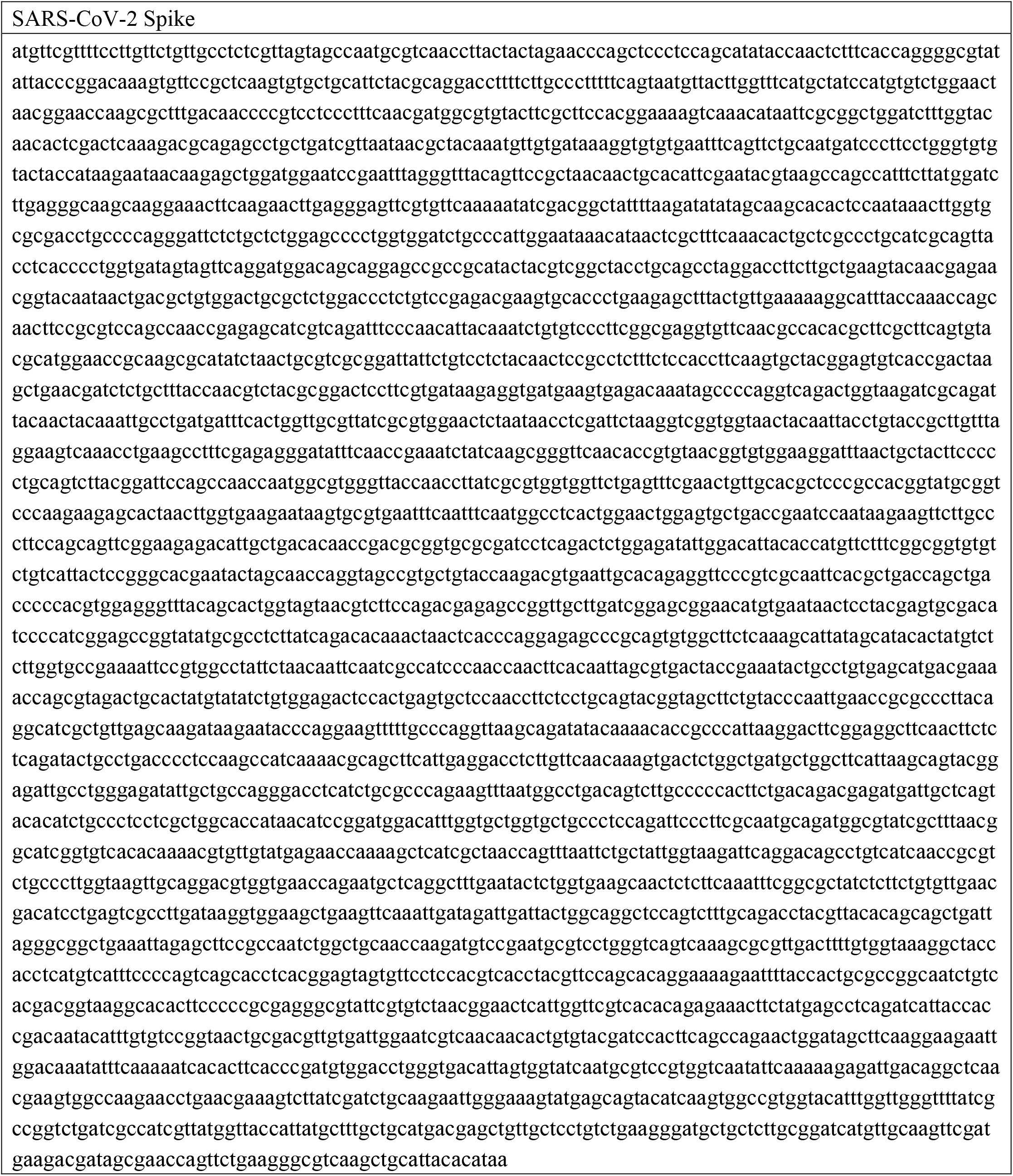
Codon-optimized DNA sequences of spike gene (without endoplasmic reticulum (ER)-retention signal)

**Supplementary Table 2.**
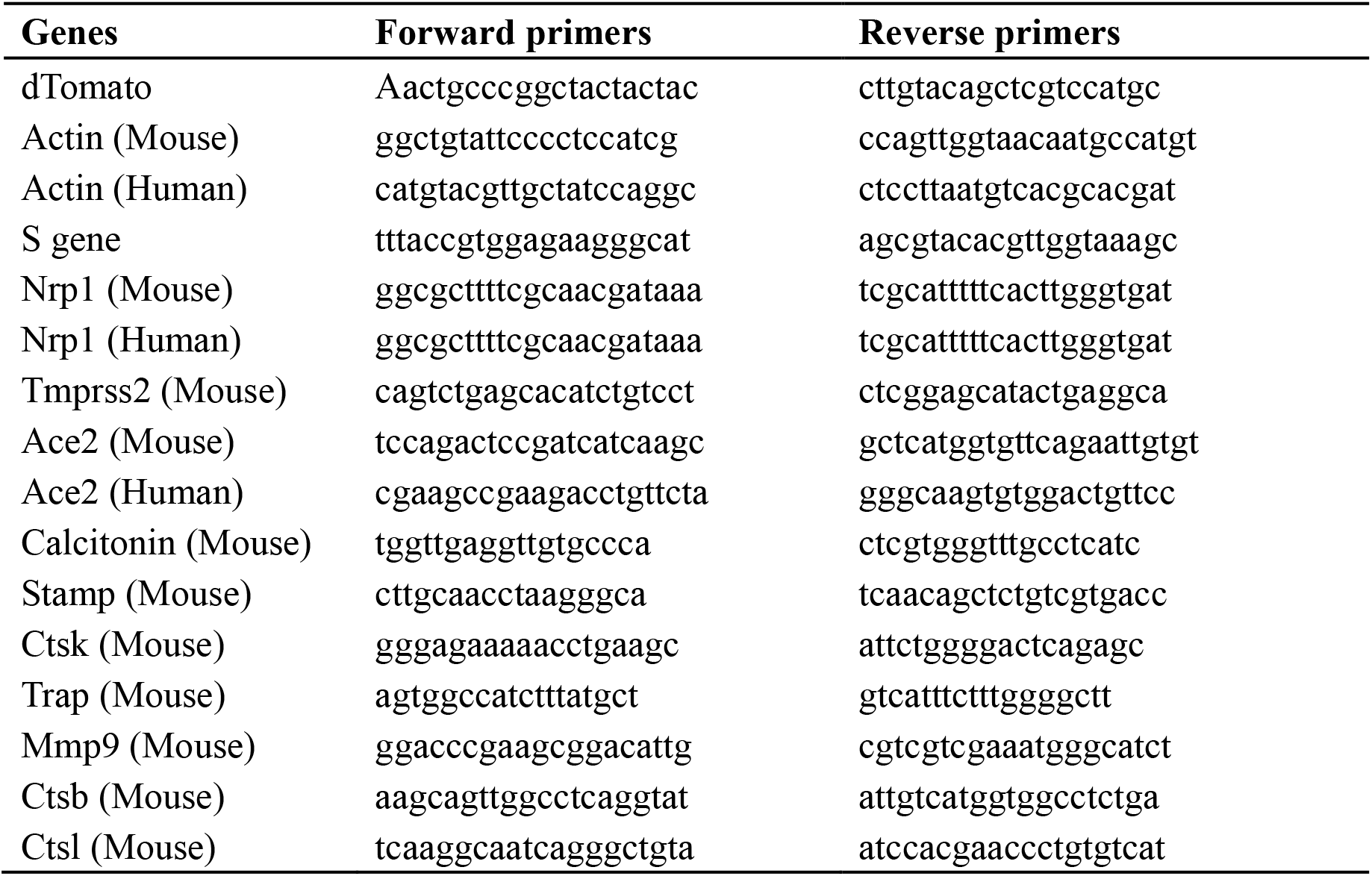
Primers for RT-qPCR.

