## Supplementary figures and images for "Neuropilin-1 Mediates SARS-CoV-2 Infection in Bone Marrow-derived Macrophages"

### Extended Data Fig 4 Single cell transcriptome analysis of brain or bone marrow cells directly isolated from 1 --, 6 and 20 month mice.

**Extended Data Figure4**

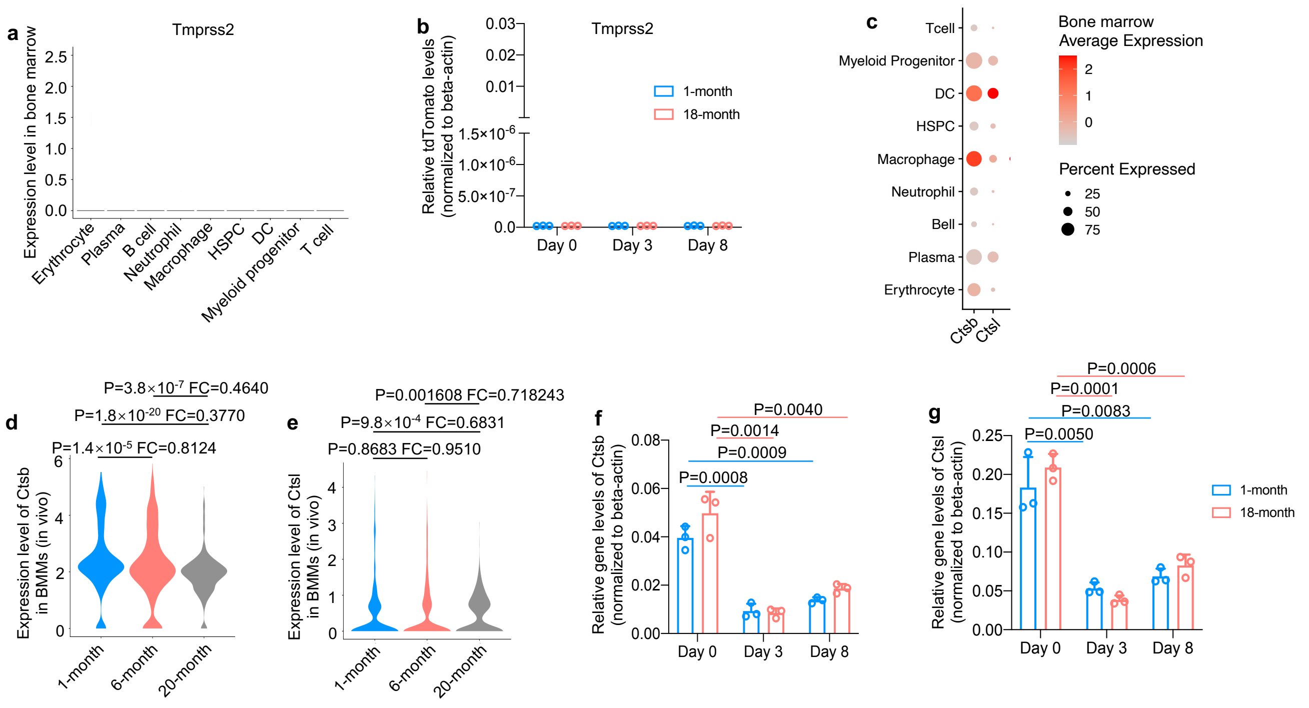

### Extended Data Fig. 1 Single cell transcriptome analysis of SARS CoV 2 pseudovirus infection in cultured BMMs from 1 and 18 month mice. a

**Extended Data Figure1**

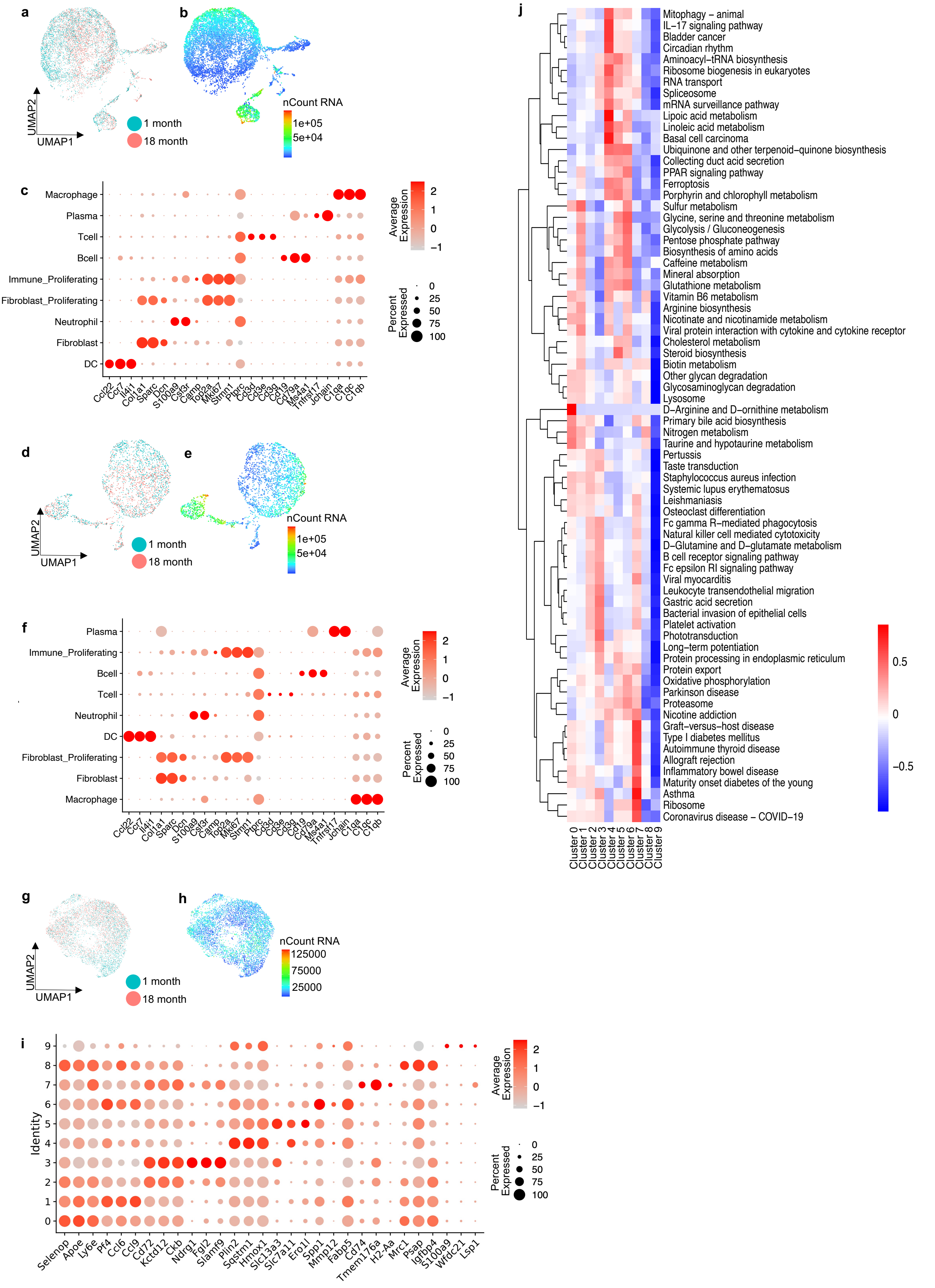

### Extended Data Fig. 2 Single cell transcriptome analysis of BMMs and brain directly isolated from 1 --, 6 and 20 month mice.

##### Extended Data Figure2

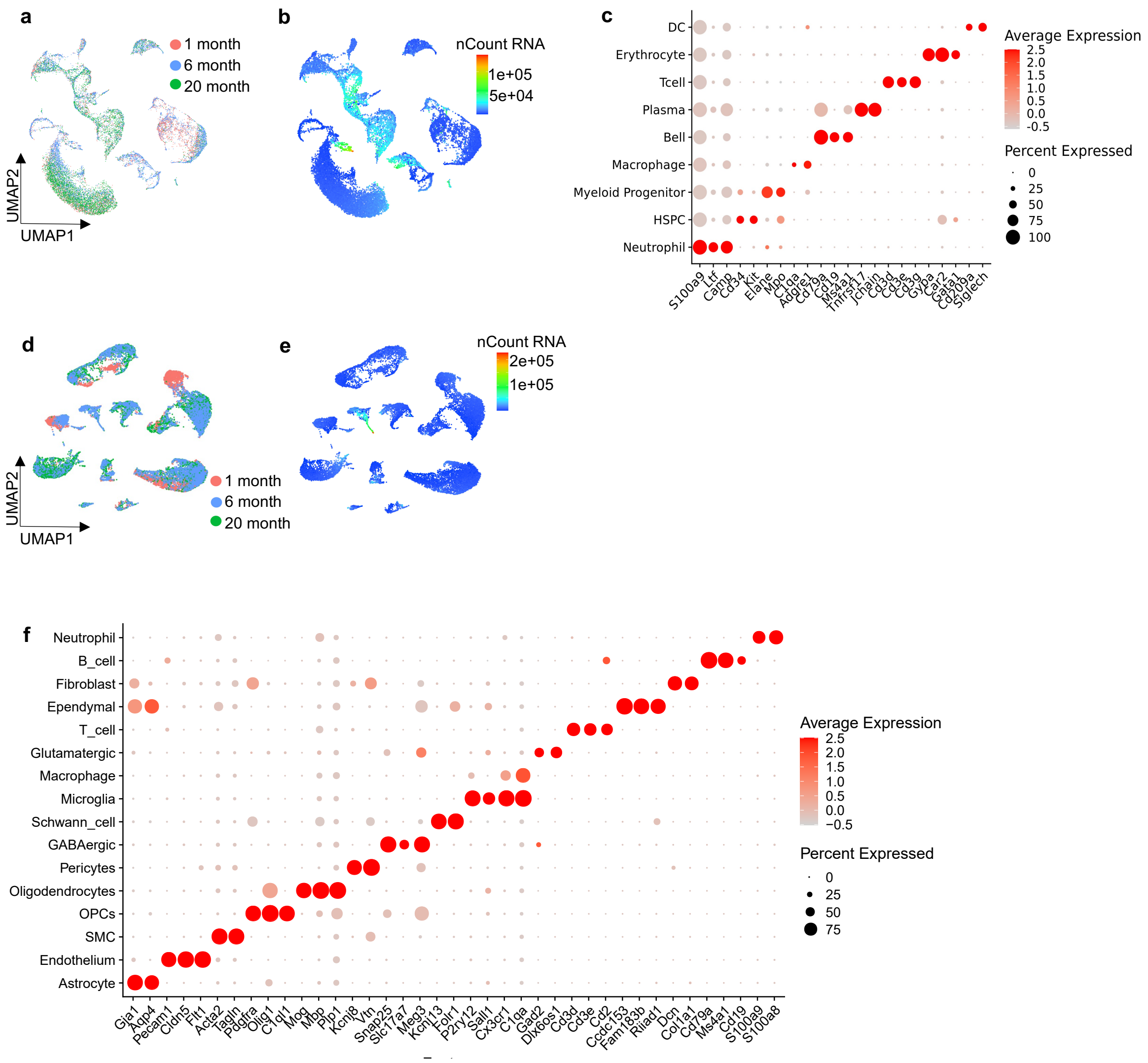

### Extended Data Fig. 3 RT qPCR quantification of altered gene expression during BMM to osteoclast differentiation.

#### Extended Data Figure3

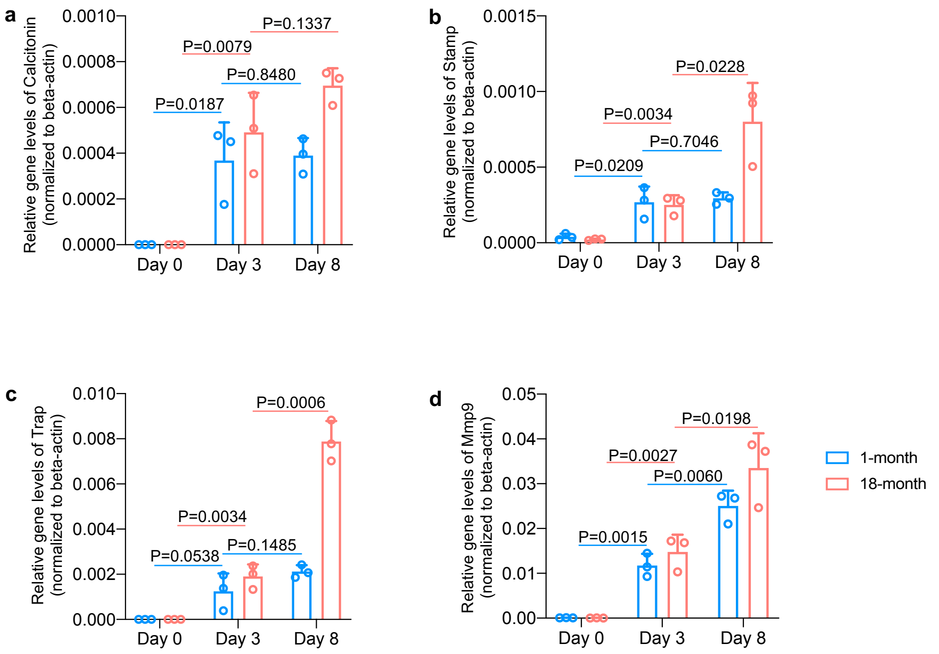
